# Identification of substrates for the conserved prolyl hydroxylase Ofd1 using quantitative proteomics in fission yeast

**DOI:** 10.1101/363747

**Authors:** He Gu, Bridget T. Hughes, Peter J. Espenshade

## Abstract

Prolyl hydroxylation functions in diverse cellular pathways, such as collagen biogenesis, oxygen sensing, and translation termination. Prolyl hydroxylation is catalyzed by 2-oxoglutarate (2-OG) oxygenases. The fission yeast 2-OG oxygenase Ofd1 dihydroxylates the 40S ribosomal protein Rps23 and regulates the hypoxic response by controlling activity and stability of the sterol regulatory element-binding protein Sre1. Multiple substrates have been found for 2-OG oxygenases, yet the only known substrate of Ofd1 and its homologs is Rps23. Here, we report the first fission yeast prolyl hydroxylome and demonstrate that hydroxylation is more prevalent than previously known. Using quantitative mass spectrometry, we identify Rpb10, a shared subunit in RNA polymerase I, II, and III, as a novel Ofd1 substrate. In addition, we discovered six Ofd1 binding partners and 16 additional Ofd1 candidate substrates. Although Ofd1 promotes Sre1 degradation, proteomic analysis revealed that Ofd1 does not broadly regulate protein degradation. Instead, the effect of Ofd1 on the proteome is through negative regulation of Sre1N. Finally, we show that the interaction between Ofd1 and the Sre1 bHLH region is conserved across Sre1 homologs suggesting that Ofd1-dependent regulation of SREBPs may be conserved in other fungi. Collectively, these studies provide a new dataset of post-translational modifications and expand the biological functions for a conserved prolyl hydroxylase.

## Introduction

Eukaryotes require environmental oxygen for essential metabolic processes. Cells exist in environments with varying oxygen supply and thus initiate biological responses to maintain homeostasis. Prolyl hydroxylation is an oxygen-dependent post-translational modification that is modulated in a variety of fundamental biological processes and diseases, such as cancer [1-3]. The modification is catalyzed by prolyl hydroxylases, whose activity depend on the Kreb’s cycle intermediate 2-oxoglutarate (2-OG), Fe(II) and oxygen [4]. The addition of a hydroxyl group on the proline residue affects protein interactions, and the oxygen requirement for this post-translational modification allows oxygen supply to regulate signaling [5]. One well-studied mechanism involving prolyl hydroxylation is regulation of the oxygen-responsive hypoxia-inducible transcription factor 1 alpha (HIF-1α). Under normoxia, hydroxylation of HIF-1α by HIF prolyl hydroxylase domain proteins (PHDs) recruit the von Hippel-Lindau (pVHL) E3 ubiquitin ligase, leading to HIF-1α ubiquitination and proteasomal degradation. Under hypoxia, the PHD substrate oxygen is limiting. HIF-1α accumulates and binds HIF-1β to activate transcription of genes that are involved in the hypoxic response and crucial aspects of cancer biology [6, 7].

Our discovery that the fission yeast sterol regulatory element-binding protein (SREBP) Sre1 is an oxygen-regulated transcription factor established a new mechanism for hypoxic signal transduction [8]. Sre1 is an ER membrane protein that is transported to the Golgi where its N-terminal transcription factor (Sre1N) is released in response to a reduction in oxygen or ergosterol [8]. Sre1 and Sre1N-like proteins are conserved across fungi, where they have been shown to regulate sterol synthesis and the hypoxic response [9-12]. Ofd1, a 2-OG/Fe(II)-dependent oxygenase, regulates the stability and activity of Sre1N through oxygen-regulated protein-protein interactions [13]. Binding of Ofd1 to Sre1N prevents Sre1N from binding to DNA and leads to Sre1N proteasomal degradation [14, 15]. Under hypoxia, Ofd1 is sequestered by the ribosomal protein Rps23 and the nuclear adaptor Nro1, allowing Sre1N to accumulate and activate hypoxic genes transcription [16, 17]. In addition, Ofd1 hydroxylates Rps23 Pro62 [17, 18]. Ofd1 is conserved in budding yeast (Tpa1), *Drosophila* (Sud1) and humans (OGFOD1), where Rps23 hydroxylation is important for translational fidelity [18-20]. OGFOD1 catalyzes prolyl trans-3-hydroxylation whereas the fungal enzymes Ofd1 and Tpa1 catalyze prolyl 3,4-dihydroxylation, a novel post-translational modification [18].

Prolyl hydroxylation was first identified in collagen biosynthesis in 1967 [21, 22]. In the past two decades, emerging functional studies suggest prolyl hydroxylation functions not only in the hypoxic response, but also in diverse cellular pathways [1, 5, 23]. For example, in addition to the prolyl hydroxylation of HIF-1α, PHD3 also catalyzes the prolyl hydroxylation of β_2_-adrenergic receptor, erythropoietin receptor, pyruvate kinase 2, and ATF-4 [24-27]. Despite these advances, Rps23 remains the only known substrate for Ofd1 and its human homolog OGFOD1. In this study, we identify additional Ofd1 substrates using quantitative mass spectrometry analysis and report the first fission yeast prolyl hydroxylome. Our analysis also revealed that the principal effect of Ofd1 on the proteome is through negative regulation of Sre1-dependent transcription, which may be a conserved regulatory mechanism in fungi.

## Materials and methods

### Materials

Yeast extract was obtained from Becton Dickinson and Co.; amino acids for the medium from Sigma; Edinburgh minimal medium from MP Biomedical. Oligonucleotides were provided by Integrated DNA Technologies. Heavy lysine (^13^C_6_/^15^N_2_) was purchased from Cambridge Isotope Laboratories. Common lab reagents were obtained from either Sigma or ThermoFisher Scientific.

### Yeast strains and culture

Wild-type haploid *S. pombe* KGY425 (*h*-, *his3*-*D1*, *leu1*-*32*, *ura4*-*D18*, *ade6*-*M210*) was obtained from American Type Culture Collection (Burke and Gould, 1994). *S. pombe* strain *ofd1Δ* (PEY1801) has been described earlier [13, 17]. *S. pombe eIF3m*-*TAP* (HGY10) and *ofd1Δ eIF3m*-*TAP* (HGY11) were generated from haploid yeast (KGY425, PEY2801) by homologous recombination using established techniques with pFA6a-CTAP-MX6 plasmid [13, 28].

Yeast strains were grown to exponential phase at 30°C in yeast extract plus supplements (225 μg/ml each of histidine, leucine, adenine, lysine and uracil) or in SILAC medium (Edinburgh minimal medium plus 75 μg/ml each of histidine, adenine, leucine and uracil and 30 μg/ml of regular or heavy lysine) [29].

### Yeast two-hybrid screen

Pairwise yeast two-hybrid assays were performed following the Clontech Matchmaker™ GAL4 Two-Hybrid System 3 user manual, using the yeast two-hybrid *S. cerevisiae* strain AH109 [30]. Full length Ofd1 was cloned into pGBKT7 (bait plasmid) as described previously [17]. For the screen of Ofd1 binding partners, *Schizosaccharomyces pombe* cDNA Matchmaker library (XL4000AA, Clontech) was used as prey. Prey and bait plasmids were co-transformed into AH109 competent cells. In total, we screened 3.1 × 10^5^ transformants. Plasmid DNA was isolated from positive colonies and sequenced. In-frame cDNA fragments corresponding to 40 proteins were identified in the screen. Their full-length cDNAs were cloned to pGADT7 and assayed for Ofd1 binding in the same yeast two-hybrid system. Equal number of transformed cells was plated on control plate (DDO, SD –Leu –Trp) and selective plate (QDO, SD –Leu –Trp –Ade – His, supplemented with X-α-Gal). Plates were incubated at 30°C for 4 days before imaging with a flatbed scanner.

For the screen of bHLH domains of SREBP and SREBP-N proteins, DNA fragments encoding the bHLH domains of selective SREBP and SREBP-N proteins were synthesized by Integrated DNA Technologies and cloned into pGADT7 (prey plasmid). The yeast two-hybrid assay was performed the same way as described above. *S. cerevisiae* colonies on control plate were inoculated and cultured in SD - Leu - Trp liquid medium for protein expression test. Yeast were grown to OD_600_ ~ 0.7 and lysates were prepared as described previously [8]. Gal4-AD-bHLH fusion proteins were resolved by SDS-PAGE and detected by immunoblotting using the anti-HA 12CA5 monoclonal antibody (1:1000) (Sigma-Aldrich) as previously described [8, 31].

### TMT mass spectrometry analysis of *S. pombe* proteomics

#### Sample preparation

Five biological replicates of *S. pombe WT* and *ofd1Δ* cells were cultured in 400 ml EMM medium (Edinburgh minimal medium plus 75 mg/l leucine, histidine, adenine, uracil, and 30 mg/l lysine) to OD_600_ ~ 0.7. Cells were harvested by spinning at 4,000 xg for 5 min. Cell pellets were resuspended in 10 ml cold lysis buffer (50 mM Tris-HCl pH 7.5, 150 mM NaCl, 1x EDTA-free Protease Inhibitor Cocktail) before lysing using an Avestin Emulsiflex C3 high pressure homogenizer. NP-40 was added to the crude lysate to 0.1% (v/v) final concentration and rotated at 4ºC for 30 min. The crude lysate was centrifuged at 20,000 xg for 30 min. The supernatant was collected and protein concentration determined by Pierce BCA Protein Assay (Thermo Fisher Scientific). Proteins (200 μg) from each of the 10 samples were TCA precipitated and washed by acetone twice.

Precipitated protein extracts (200 μg for each sample) were re-solubilized in 200 μl of 100 mM triethyl ammonium bicarbonate (TEAB) pH 8.5 and reduced with 10 μl of 200 mM TCEP then alkylated with 10 μl of 375 mM iodoacetamide in the dark for 30 min. Reduced and alkylated proteins were digested overnight at 37ºC by adding 10 μg of Trypsin/LysC mixture (V5071, Promega) in 100 mM TEAB. The next morning, 10 μl of 10% (v/v) TFA was added and samples were evaporated to dryness in a speedvac. Peptides were re-solubilized in 200 μl of 100 mM TEAB and divided into four 50 μg aliquots.

#### TMT-labeling and fractionation

Individual samples (50 μg) were labeled with a unique isobaric mass tag reagent (TMT 10-plex, Thermo) according to the manufacturer’s instructions. Both pairing and labeling order of TMT reagent and peptide sample were randomized. Briefly, TMT-10plex reagents (0.8 mg vials) were allowed to come to room temperature before adding 41 μl of anhydrous acetonitrile, then briefly vortexed and centrifuged. The entire TMT reagent vial was added to the 50 μg peptide sample and reacted at room temperature for 1 hr. 5% (v/v) hydroxylamine (8 μl) was then added to quench the reaction. The 10 TMT-labeled samples were then combined and vacuum centrifuged to dryness. The combined sample of TMT-labeled peptides was resuspended in 2 ml of 10 mM TEAB and separated into 84 fractions at 250 μl/min using a 0-90% acetonitrile gradient in 10 mM TEAB on a 150 mm x 2.1 mm ID Waters XBridge 5 μm C18 using an Agilent 1200 capillary HPLC in normal flow mode and Agilent 1260 micro-fraction collector. The 84 fractions were concatenated into 24 fractions by combining all odd rows of each column 1 through 12 into 12 fractions and all even rows of each column into another 12 fractions.

#### LC-MS/MS

An aliquot (5 μl) of each of the 24 fractions was analyzed by reverse phase liquid chromatography coupled to tandem mass spectrometry. Briefly, peptides were separated on a 75 μm x 150 mm in-house packed Magic C18AQ column (5 μm, 120 Å (Bischoff) using 2-90% acetonitrile gradient in 0.1% (v/v) formic acid at 300 nl/min over 90 min on a EasyLC nanoLC 1000 (Thermo). Eluting peptides were sprayed through 1 μm emitter tip (New Objective) at 2.0 kV directly into the Q-Exactive. Survey scans (full ms) were acquired from 350-1700 m/z with data dependent monitoring of up to 15 peptide masses (precursor ions), each individually isolated in a 1.5 Da window and fragmented using HCD (higher-energy collisional dissociation) with a normalized collision energy (NCE) of 30 and a 30 second dynamic exclusion. Precursor and fragment ions were analyzed at resolutions 70,000 and 35,000, respectively, with automatic gain control (AGC) target values at 3e6 with 100 ms maximum injection time (IT) and 1e5 with 250 ms maximum IT, respectively.

#### Database search

MS data were searched against the *S. pombe* RefSeq2014 database in Proteome Discoverer (version 1.4, Thermo Fisher Scientific) using Mascot alone and Mascot run through the MS2-processor node. All peptides were searched with a 20 ppm tolerance MS and 0.03 for MS2 and filtered at a 1% FDR. Dynamic modification was chosen for carbamidomethyl C, oxidation M, deamidation of N and Q, hydroxylation of P, dihydroxylation of P, TMT 6-plex for N-term and K.

#### Data processing

The protein area calculated by Proteome Discoverer Software was used to quantify protein abundance. The relative abundance of each protein in each of the 10 TMT channels was calculated as following:

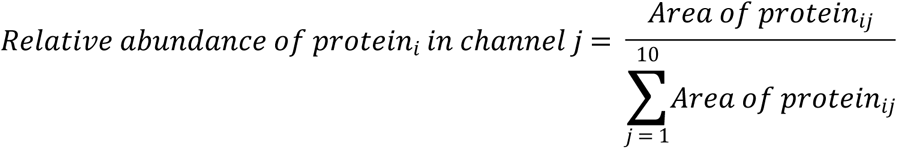

The relative abundance of *WT*5 and *ofd1Δ*2 samples were higher than the other replicates and therefore these two samples were excluded from quantile normalization and further analysis. The remaining eight samples were quantile normalized using ‘preprocessCore’ package embedded in R [32]. The normalized relative abundance was saved and used for further analysis. Student’s t-test was applied to calculate the p-value of protein relative abundance in each sample using R.

At the peptide level, the precursor area was used to quantify PSM abundance. The relative abundance of each PSM in each of the 8 channels was calculated as following:

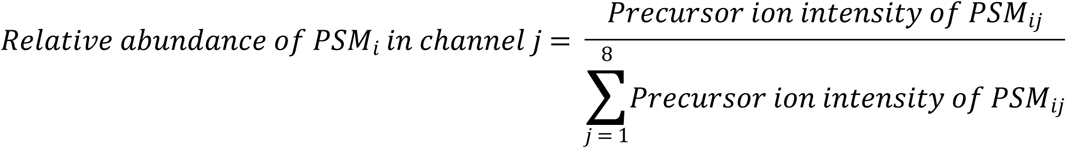

We searched for prolyl-hydroxylated PSMs that have +16 Da or +32 Da mass shifts on proline residue and have complete y- or b-series ions around the hydroxylated proline. Fold changes were calculated by the ratio of the average relative abundance in four *WT* replicates and the average relative abundance of four *ofd1Δ* replicates.

### SILAC mass spectrometry analysis of eIF3m prep

#### SILAC culture and sample preparation

HGY10 (*h*-, *his3*-*D1*, *leu1*-*32*, *ura4*-*D18*, *ade6*-*M210*, *eIF3m*-*TAP*-*KanMX6*) and HGY11 (*h*-, *his3*-*D1*, *leu1*-*32*, *ura4*-*D18*, *ade6*-*M210*, *ofd1Δ::natMX6*, *eIF3m*-*TAP*-*KanMX6*) were cultured for 15-18 generations in 1.5 L SILAC medium (Edinburgh minimal medium plus 75 mg/l leucine, histidine, adenine, uracil, and 30 mg/l heavy (HGY10) or light (HGY11) lysine) to OD_600_ ~ 0.7 as previously described [29]. Cells were harvested by spinning at 4,000 xg for 12 min. Cell pellets were resuspended in 10 ml cold lysis buffer (50 mM Tris-HCl pH 7.5, 150 mM NaCl, 1x EDTA-free Protease Inhibitor Cocktail) before lysing using Avestin Emulsiflex C3 high pressure homogenizer. NP-40 was added to the crude lysate to 0.1% (v/v) final concentration and rotated at 4ºC for 30 min. The crude lysate was centrifuged at 20,000 xg for 30 min. The supernatant was collected, and protein concentration was measured by Pierce BCA Protein Assay (Thermo Fisher Scientific). A sample of heavy lysate (300 μg) was saved for lysine incorporation analysis. Equal amounts of heavy and light lysates (1275 mg lysate each) were mixed, and eIF3m prep was purified by Protein A and calmodulin binding peptide affinity purification in sequence as described previously [33]. eIF3m-interacting proteins (54 μg) were TCA precipitated and analyzed by SILAC mass spectrometry.

eIF3m prep sample was brought up in 100 μl of 20 mM ammonium bicarbonate pH8.5 then reduced with 50 mM DTT and alkylated with 50 mM iodoacetamide. Sample was digested overnight at 37ºC with 2 μg LysC (Promega). An additional 1 μg of LysC was added the next morning for another 4-hr digestion. TFA (10 μl of 10% v/v) was added to stop the digestion. The sample was evaporated to dryness and then brought up in 2 ml of 10 mM TEAB.

#### Fractionation

Fractionation was performed in the same way as described in the TMT mass spectrometry section, except that the sample (54 μg) was used for fractionation here.

#### LC-MS/MS

The LC-MS/MS was performed in the same way as described in the TMT mass spectrometry section, except that the collision energy was 27 and the fragment ions were analyzed at a resolution of 17500.

#### Database search

Data was searched against the *S. pombe* RefSeq2014 database as described in the TMT mass spectrometry section. Note that for Ofd1 candidate substrates 10% FDR was used as indicated in the results section. Dynamic modification was chosen for carbamidomethyl C, oxidation M, deamidation of N and Q, hydroxylation of P, dihydroxylation of P, and Label: 13C(6) 15N(2) for heavy lysine (K+8.014).

#### Data processing

*S. pombe* is a lysine prototrophic microorganism. Although lysine supplement reduces lysine biosynthesis, the heavy lysine uptake in nSILAC method is always incomplete [29]. Therefore, it is important to normalize the incorporation level of the heavy lysine before comparing heavy to light ratios (H/L). To calculate the incorporation level, we analyzed the whole cell lysate of the *WT* cell (cultured in heavy SILAC medium). The heavy lysine incorporation rate for each PSM was calculated using heavy to light ratio (H/L) of heavy (HGY10) lysate.

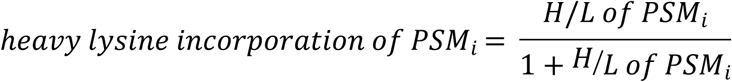

The median of heavy lysine incorporation was used for SILAC sample H/L normalization.

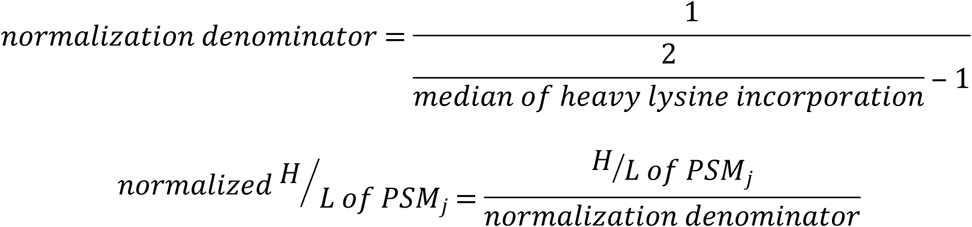

The normalized H/L was saved and used for further analysis. We searched for prolyl-hydroxylated PSMs that have +16 Da or +32 Da mass shifts on proline residues and plotted against precursor ion intensity in R. Prolyl hydroxylated PSMs with H/L > 3.5 were defined as Ofd1-dependent, and PSMs with H/L ≤ 3.5 were defined as Ofd1-independent. y- and b-series ions of each PSM were then manually inspected. The Ofd1-independent prolyl hydroxylated PSMs with complete y- and b-series ions were reported. The Ofd1-dependent prolyl hydroxylated PSMs and the assumptions about hydroxylation residues were reported.

### Multiple sequence alignment

Protein sequences were downloaded from Pombase (for *S. pombe* proteins) and NCBI database (for proteins from other species). Alignments were performed using T-coffee [34, 35], and fastaaln results were saved. Alignment figures were generated using ESPript 3 and Tpa1 crystal structure (PDB: 3KT1) was used to estimate secondary structure of Ofd1 homologs [36, 37].

## Results

### Ofd1 binding partners are candidate substrates for prolyl hydroxylation

To date, the only known enzymatic substrate for Ofd1 is Rps23 [17, 18]. However, many 2-OG dioxygenases have more than one substrate [1]. To identify additional substrates of Ofd1, we performed a yeast two-hybrid screen with *S. pombe* Ofd1 as bait and a *S. pombe* cDNA library as prey. The screen revealed six new candidate Ofd1 binding partners and confirmed the known interaction with Rps23 (Fig 1) [17, 18]. Interestingly, the screen identified two subunits of the eukaryotic translation initiation factor (eIF3) complex, Tif35 (eIF3g) and Sum1 (eIF3i). Next, we examined if eIF3 subunits were substrates of Ofd1.

**Fig 1.**
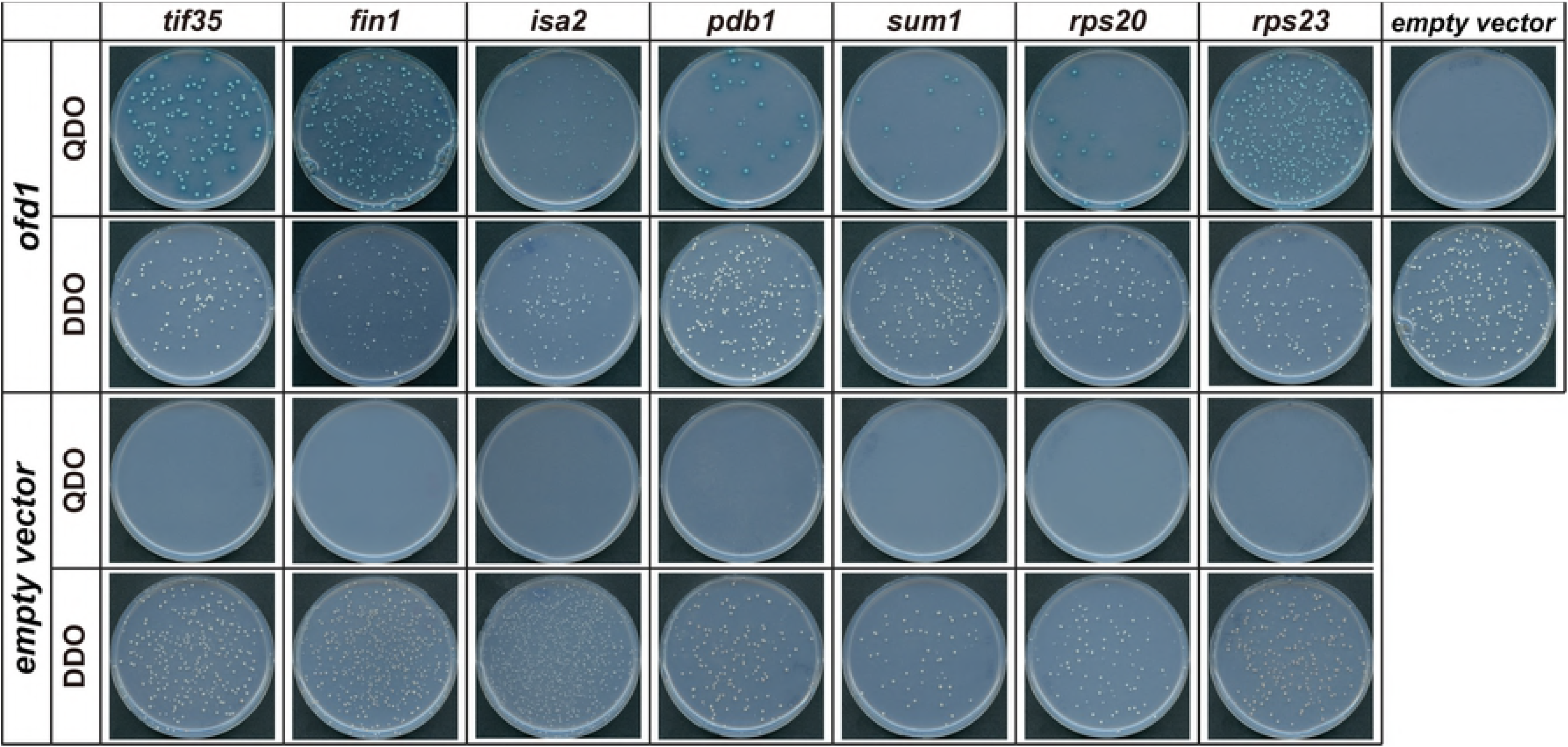
Ofd1 binding partners. Yeast two-hybrid interaction between Ofd1 and candidate binding partners. Ofd1 was fused with GAL4 DNA binding domain and *S. pombe* cDNA was fused with GAL4 activation domain. DDO: control plate, SD –Leu –Trp. QDO: selection plate, SD –Leu –Trp –Ade –His, supplemented with X-α-Gal.

### Prolyl hydroxylation in the eIF3m prep

Fission yeast has two distinct eIF3 complexes that share 6 subunits (eIF3a, b, c, f, g, i) and are distinguished by the presence of either eIF3e or eIF3m [38, 39]. The two eIF3 complexes associate with different sets of mRNAs, and the eIF3m complex contains more subunits than the eIF3e complex [38]. We tagged eIF3m with a C-terminal TAP tag to capture the largest number of eIF3 proteins and employed native stable isotope labeling by amino acids in cell culture (nSILAC) to test whether purified eIF3g and eIF3i showed Ofd1-dependent prolyl hydroxylation [29]. Wild-type (*WT*) *S. pombe* cells were labeled with heavy lysine 8 (~80% efficiency, Fig S1A), and *ofd1Δ* cells were labeled with light lysine (Fig 2A). Equal amounts of *WT* and *ofd1*Δ lysates were mixed, and eIF3m interacting proteins were purified using sequential Protein A and calmodulin binding peptide affinity purification [40, 41] (Fig 2B).

**Fig 2.**
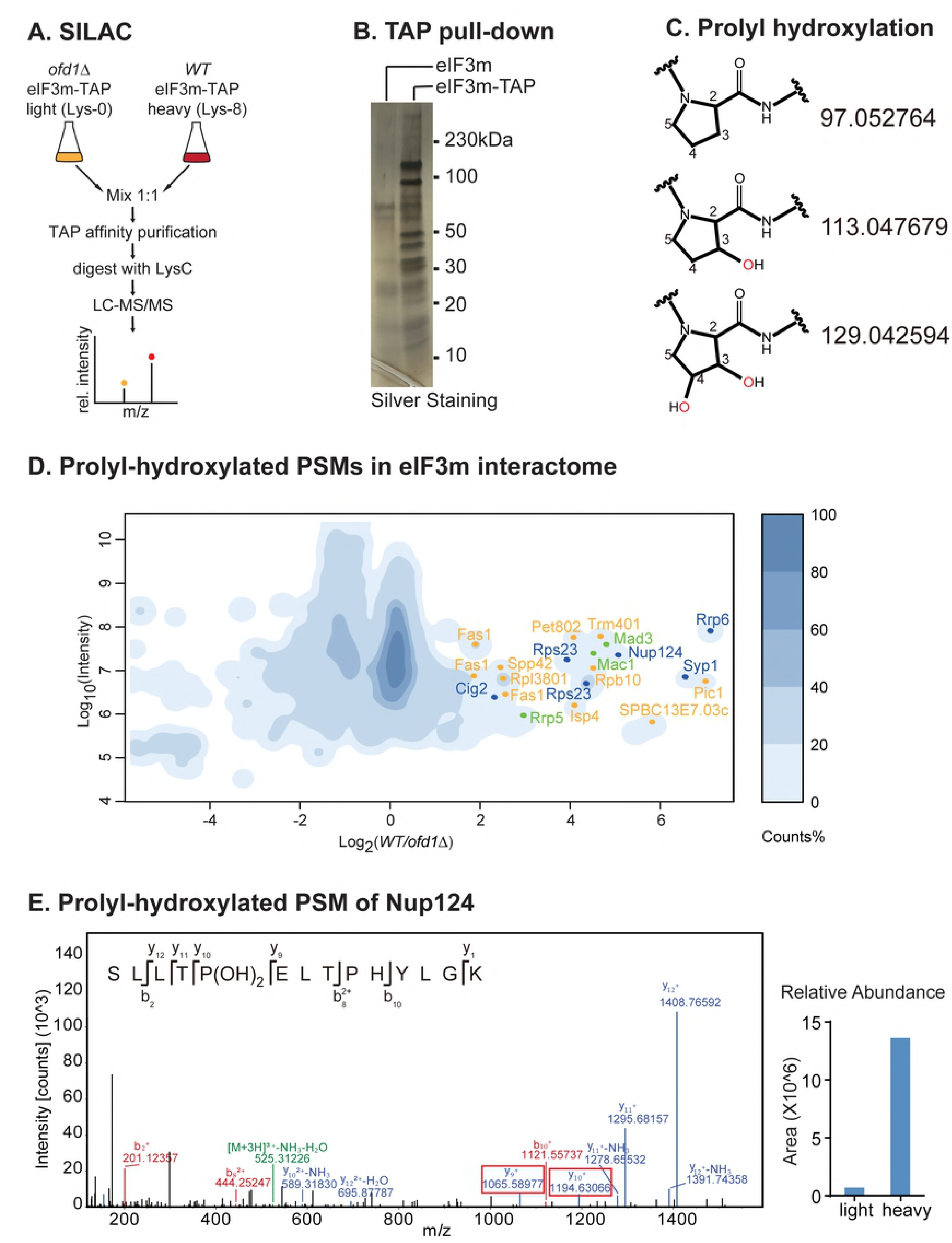
Ofd1 candidate prolyl hydroxylated substrates. (A) Design of SILAC mass spectrometry experiment comparing eIF3m prep from wild-type (*WT*) and *ofd1*Δ cells. (B) Silver stained gel of eIF3m prep affinity-purified using eIF3m-TAP. (C) Structure and mono-isotopic mass of non-hydroxylated, mono-hydroxylated and di-hydroxylated proline residues. (D) Prolyl hydroxylated peptides identified from eIF3m prep. Each prolyl-hydroxylated peptide spectrum match (PSM) was plotted. The relative abundance of each peptide in *WT* and *ofd1*Δ cells was quantified by SILAC. PSMs with heavy to light ratio greater than 3.5 were classified as Ofd1 candidate substrates. Assumptions about hydroxylation of other residues were made for the PSMs without complete b- or y-series (see Table 1). Ofd1 candidate substrates were color coded: yellow, monohydroxylated; blue: dihydroxylated; and green: one monohydroxylated residue and one dihydroxylated residue. (E) Spectrum of prolyl dihydroxylated peptide in Nup124.

Mass spectrometry analysis identified all known eIF3m complex subunits (eIF3a-d, eIF3f-i, eIF3m) with coverage ranging from 13.25% to 63.12%. This experiment also confirmed that eIF3e is not present in the eIF3m complex. We searched for peptides with +16 Da and +32 Da shifts on proline residues, corresponding to mono and dihydroxylation, respectively (Fig 2C). In the eIF3m complex, we identified 117 peptides, 43 (36.7%) of which were proline-containing peptides. One peptide from Tif32 (eIF3a) contained a monohydroxylated proline residue with 0.340 heavy to light ratio (wild-type/*ofd1Δ* cells), indicating that the modification is Ofd1-independent (Fig S3). Thus, while the yeast two-hybrid screen indicates that Ofd1 binds to eIF3 subunits, we did not find evidence that eIF3 subunits are Ofd1 substrates.

In addition to subunits of the eIF3m complex, we identified 1263 proteins in the eIF3m prep, including eIF3-interacting proteins and abundant cytosolic proteins, comparable to mass spectrometry searches in previously published results [39]. Using a FDR of 0.01, we identified 1523 peptides that contain proline residues, including 15 peptides with prolyl dihydroxylation, 52 peptides with prolyl monohydroxylation and one peptide with both mono and dihydroxylation. The prolyl hydroxylated PSMs corresponded to 55 proteins in the eIF3m prep.

We defined prolyl hydroxylated peptides with a heavy to light (H/L) ratio > 3.5 as Ofd1-dependent. Using this criterion, two unique prolyl dihydroxylated peptides from Rps23 (IGVEAKQP(OH)_2_NSAIRK and IGVEAKQ(deamidated)P(OH)_2_NSAIRK) had H/L ratios of 14.941 and 20.030 respectively, confirming that Rps23 P62 dihydroxylation is Ofd1-dependent and validating the dataset. To expand our search for Ofd1 substrates, we repeated the analysis using a FDR of 0.1 and plotted the prolyl hydroxylated peptides based on their H/L and precursor area (Fig 2D). Again, we defined Ofd1-dependent hydroxylated peptides as having a H/L ratio > 3.5 (Fig 2D). After manual inspection of the spectra, we identified 16 novel Ofd1 candidate substrates, including 9 monohydroxylated peptides, 4 dihydroxylated peptides, and 3 peptides containing one mono and one dihydroxylated proline (Table 1, Fig S2). The prolyl dihydroxylated peptide from Nup124, for instance, had complete y-ion series around the proline residue and was more abundant in *WT* (heavy) cells, indicating the hydroxylation was Ofd1-dependent (Fig 2E). For PSMs with incomplete b- or y-ion series, we made specific assumptions about other residue modifications in order to classify these PSMs as prolyl hydroxylated (Table 1).

**Table 1.**
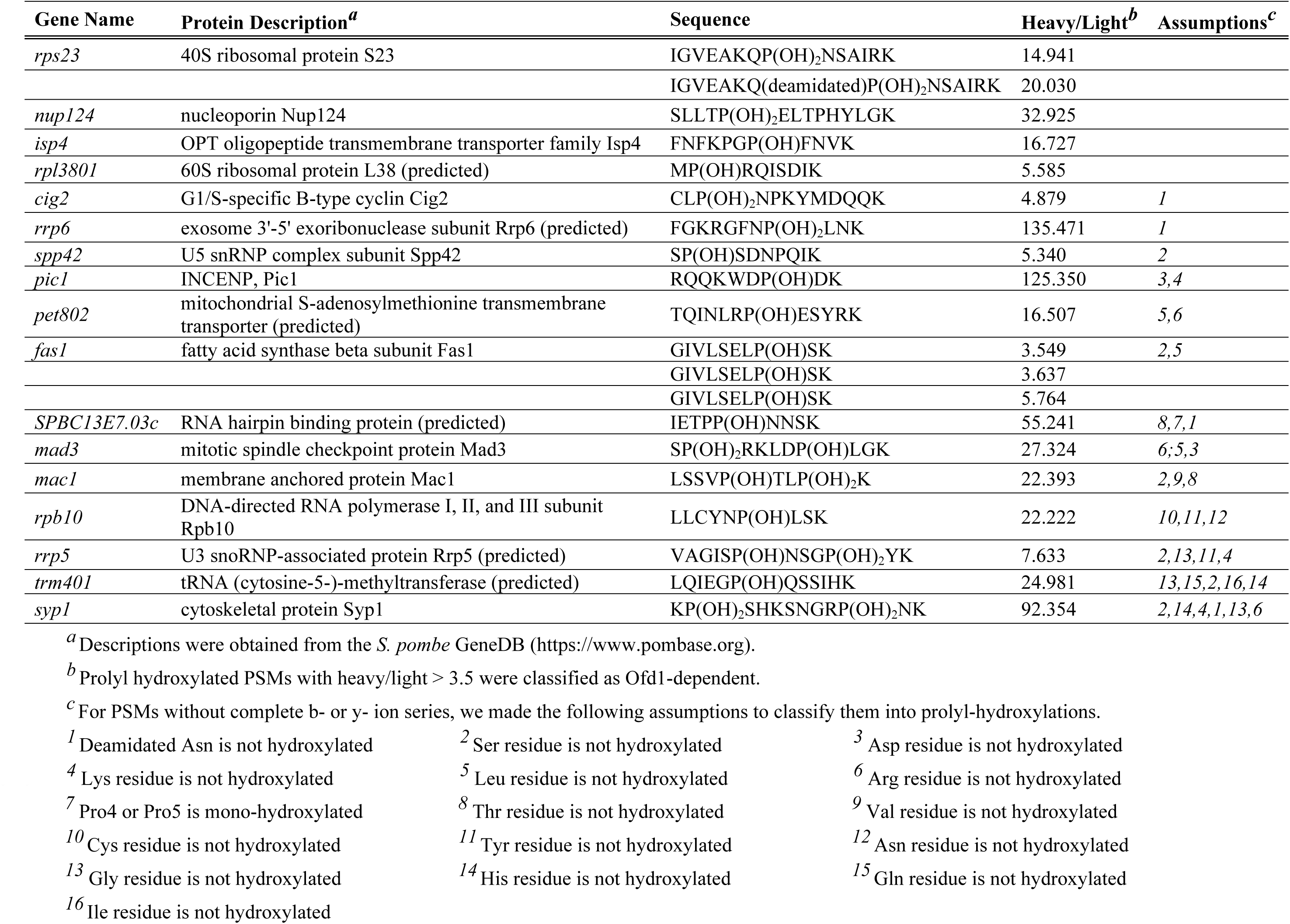
Ofd1 candidate substrates from eIF3m prep.

We recently identified homologous Ofd1 binding sequences in Rps23 and Nro1 that mediate direct binding to Ofd1 [17]. Multiple sequence alignments with the Ofd1 binding sequence in Rps23, Nro1, and Ofd1 candidate substrates revealed that a similar binding sequence exists in Pic1 (kinetochore protein, INCENP ortholog Pic1) and Fas1 (fatty acid synthase beta subunit Fas1) (Fig S1B), indicating that a similar sequence may mediate these Ofd1-substrate interactions. Together, these data indicate that additional Ofd1 substrates likely exist.

Rps23 is completely hydroxylated under normoxia [17, 18]. As expected, the averaged H/L ratio of the non-hydroxylated Rps23 P62 peptide was 0.001, indicating 100% hydroxylation, i.e. no non-hydroxylated Rps23 in *WT* cells and an accumulation of non-hydroxylated peptide in *ofd1Δ* cells. For the candidate Ofd1-dependent modifications in Table 1, we observed non-hydroxylated peptides for Fas1, Rpb10, and Rrp5. In contrast to Rps23, their corresponding H/L ratios are 0.752, 0.575, 0.570 respectively, indicating that they are not 100% hydroxylated in *WT* cells. Recognizing that prolyl-hydroxylated peptides are more hydrophilic and thus may behave differently from non-hydroxylated peptides, we compared the precursor area of hydroxylated and non-hydroxylated peptides in each yeast strain as an independent test for Ofd1-dependency. Indeed, in *WT* cells (heavy) > 99% of the peptides from Rps23 were dihydroxylated, whereas in *ofd1Δ* cells (light) 92% were non-hydroxylated (Fig S1C). The non-hydroxylated peptides for Fas1 and Rrp5 did not dramatically increase in *ofd1Δ* cells, perhaps due to the fact that the hydrophobic non-hydroxylated peptides were easier to detect. However, the non-hydroxylated peptide from Rpb10 increased from 52% in *WT* cells to 96% in *ofd1Δ* cells, indicating that monohydroxylation of Rpb10 is Ofd1-dependent (Fig S1C). Together, these data indicate that Rpb10 is an Ofd1 substrate that is present in mono-hydroxylated and non-hydroxylated forms. Additional experiments are required to quantify accurately the two Rpb10 populations.

Analysis of the complete eIF3m prep also revealed Ofd1-independent prolyl hydroxylation (H/L ≤ 3.5). Using a strict FDR of 0.01, we discovered 7 prolyl hydroxylated peptides with complete b- or y-ion series around the modified proline residue (Table 2, Fig S3). These peptides belonged to ribosomal proteins (Rps2, Rps9, Rpl10a), mitochondrial ribosomal protein subunit Rsm7, translation initiation factor eIF3a as noted above, alcohol dehydrogenase Adh1 and ubiquitin activating enzyme E1 Ptr3. In addition to Ofd1, there are 9 other predicted dioxygenases in *S. pombe* (Table 3) that may play roles in Ofd1-independent prolyl hydroxylation of these proteins. Collectively, these results suggest that prolyl hydroxylation is present in proteins isolated in our eIF3m prep.

**Table 2.**
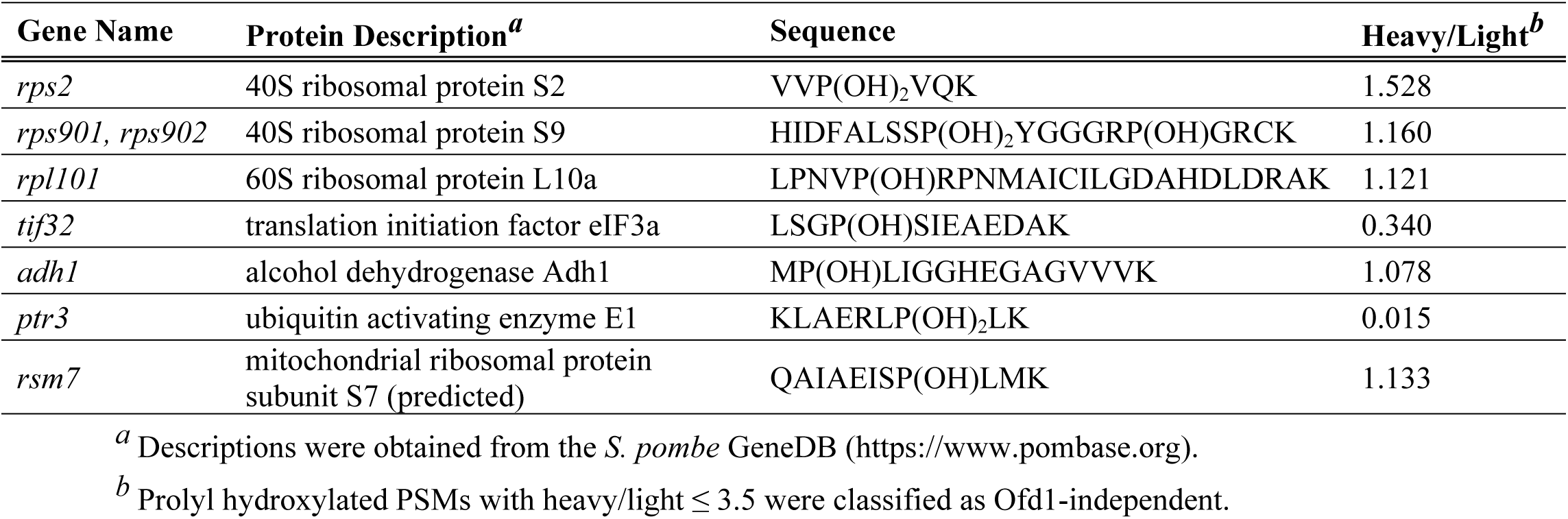
Ofd1-independent prolyl hydroxylations from eIF3m prep.

**Table 3.** Dioxygenases in *S. pombe*.

| Gene Name | Systematic ID | Protein Description<sup>a</sup> |
| --- | --- | --- |
| *ofd1* | *SPBC6B1.08c* | Prolyl 3,4-dihydroxylase Ofd1 |
| *isp7* | *SPAC25B8.13c* | 2-OG-Fe(II) oxygenase superfamily protein |
|  | *SPCC1494.01* | iron/ascorbate oxidoreductase family |
|  | *SPCC1620.13* | phosphoglycerate mutase/6-phosphofructo-2-kinase family (predicted) |
|  | *SPBC460.04c* | sulfonate/alpha-ketoglutarate dioxygenase (predicted) |
| *xan1* | *SPCC576.01c* | alpha ketoglutarate dependent xanthine dioxygenase Xan1 |
| *jmj1* | *SPAC25H1.02* | histone demethylase Jmj1 (predicted) |
| *abh1* | *SPBC13G1.04c* | tRNA demethylase (predicted) |
| *ofd2* | *SPAP8A3.02c* | histone H2A dioxygenase Ofd2 |
| *adi1* | *SPBC887.01* | acireductone dioxygenase family Adi1 (predicted) |

<sup>a</sup> Descriptions were obtained from the *S. pombe* GeneDB (https://www.pombase.org).

In the eIF3m dataset, we observed that a fraction of prolyl hydroxylated peptides (20.9%) deviated from the center with H/L ratios between 0.330 and 0.660 (Fig 2D). This phenomenon was also observed in the entire collection of peptides identified in the eIF3m prep (Fig S4A), indicating that it was independent of prolyl hydroxylation. When H/L ratios were calculated for eIF3m interacting proteins rather than peptides, the left-shifted population disappeared indicating that the phenomenon was likely due to result of noise at PSM level rather than from biological differences between *WT* and *ofd1Δ* cells (Fig S4B). In addition, the half-life of proteins containing peptides from the left-shifted population was identical to that of the entire eIF3m prep (Fig S4C), indicating the shift was not due to differences in protein turnover [42].

### Sre1 target gene products are upregulated in *ofd1*Δ cells

Our identification of novel prolyl hydroxylation in the eIF3m prep prompted us to extend our analysis to the cellular proteome. We performed quantitative proteomics analysis on whole cell lysates from *WT* and *ofd1*Δ cells employing tandem mass tag labeling and tandem mass spectrometry (LC-MS/MS) with five biological replicates for each strain. Four replicates from each strain were used for quantile normalization and further analysis (Fig S5). Overall, 2862 proteins were detected, which is comparable to published fission yeast proteomics results (Fig 3A) [42]. We searched for prolyl hydroxylation as described previously using a FDR of 0.01 and identified 38 peptides with prolyl dihydroxylation, 122 peptides with prolyl monohydroxylation, and 3 peptides with both mono and dihydroxylation. These PSMs represented a small fraction of total PSM detected (0.2%) and mapped to 128 proteins. After manual inspection, we identified 12 peptides with complete b- or y-ion series around the modified proline residue (Table 4, Fig S6). Peptide abundance was not significantly different (p-value < 0.05) between *WT* and *ofd1Δ* cells, indicating that these hydroxylations are Ofd1-independent. Overall, prolyl hydroxylated peptides represented a small fraction of total peptides detected. Development of reagents to enrich for prolyl hydroxylated peptides will improve detection of prolyl hydroxylation in proteomic samples.

**Fig 3.**
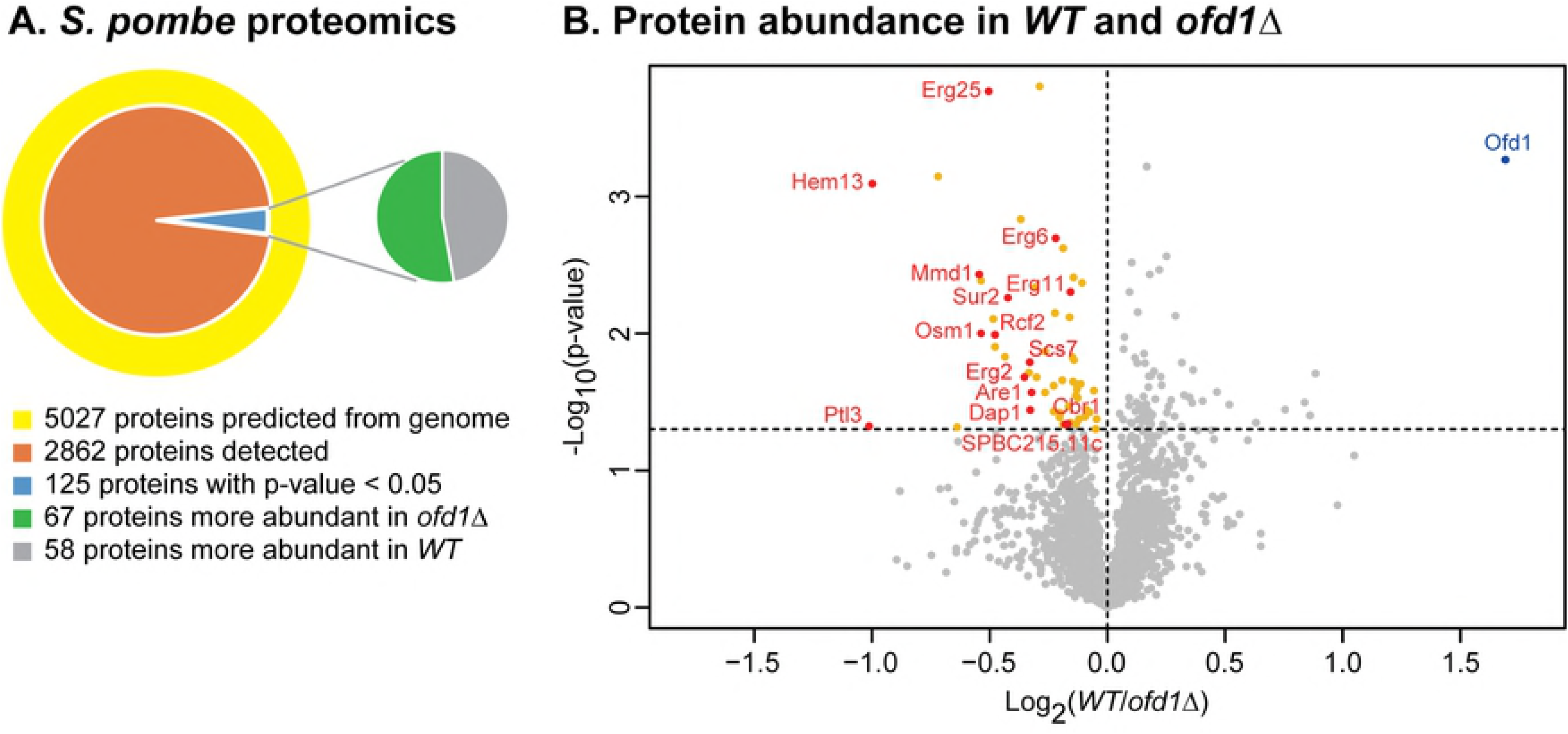
Sre1 target gene products are upregulated in *ofd1*Δ cells. Five replicates of whole cell lysates from *WT* and *ofd1Δ* cells were analyzed by tandem mass tag (TMT) labeling and tandem mass spectrometry (LC-MS/MS). For the analysis, four replicates from each strain were quantile normalized to calculate p-values and fold changes. (A) 2862 proteins of the predicted 5027 protein entries from the *S. pombe* database were identified [43]. Majority of proteins (96.7%) showed no difference (p-value > 0.05) between *WT* and *ofd1*Δ strains. (B) Proteins that accumulated significantly (p-value < 0.05) in *ofd1*Δ cells are labeled in yellow or red. Sre1 regulated gene products are labeled in red. Ofd1 is labeled in blue.

**Table 4.**
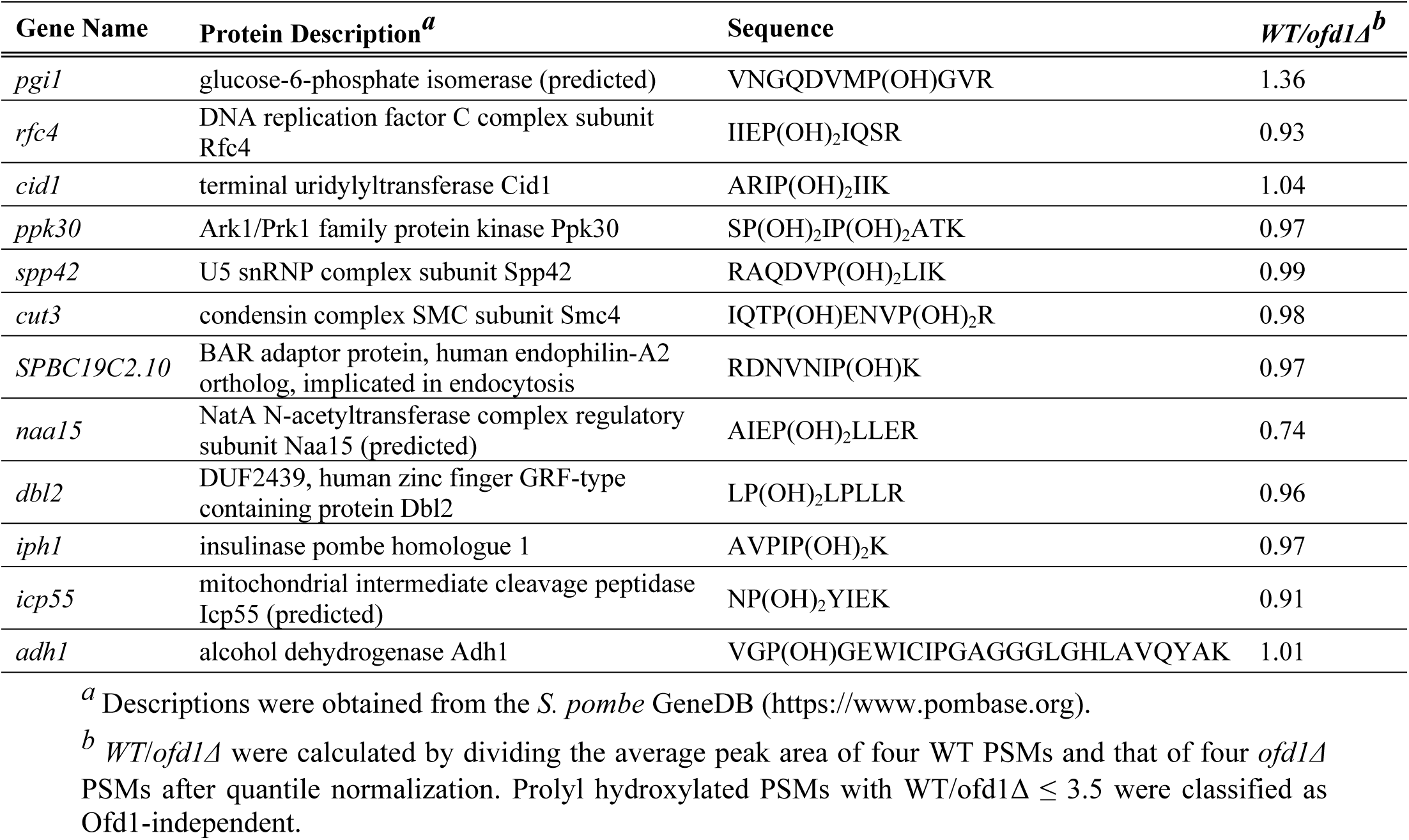
Ofd1-independent prolyl hydroxylations from *S. pombe* proteome.

At the protein level, we performed student’s t-tests to calculate the p-value of individual protein abundance between *WT* and *ofd1Δ* cells. The majority of proteins showed equal expression between *WT* and *ofd1Δ* cells. A small fraction of proteins (58, 2.0%) were significantly upregulated in *ofd1Δ* cells compared to *WT* cells, and none were upregulated greater than 2-fold. Together, these data indicate that Ofd1 does not regulate global protein expression under steady state conditions (Fig 3B). Notably, Sre1 target gene products were highly enriched in this subset of 58 proteins (Fig 3B and Table 5) [44], indicating that the principal effect of Ofd1 on the proteome is through negative regulation of Sre1-dependent transcription.

**Table 5.** Sre1 target genes with products upregulated in *ofd1Δ* cells.

| Gene Name | Systematic ID | Protein Description<sup>a</sup> |
| --- | --- | --- |
| *hem13* | *SPAC222.11* | coproporphyrinogen III oxidase Hem13 (predicted) |
| *sur2* | *SPBC887.15c* | sphingosine hydroxylase/sphingolipid delta-4 desaturase activity Sur2 |
| *osm1* | *SPAC17A2.05* | fumarate reductase Osm1 (predicted) |
| *mmd1* | *SPAC30C2.02* | deoxyhypusine hydroxylase |
| *erg2* | *SPAC20G8.07c* | C-8 sterol isomerase Erg2 |
| *erg6* | *SPBC16E9.05* | sterol 24-C-methyltransferase Erg6 |
| *erg11* | *SPAC13A11.02c* | sterol 14-demethylase Erg11 |
| *erg25* | *SPAC630.08c* | C-4 methylsterol oxidase Erg25 (predicted) |
| *rcf2* | *SPAC1565.01* | cytochrome c oxidase assembly protein Rcf2 |
| *are1* | *SPAC13G7.05* | acyl-CoA-sterol acyltransferase Are1 (predicted) |
| *scs7* | *SPAC19G12.08* | ER sphingosine hydroxylase Scs7 |
| *dap1* | *SPAC25B8.01* | cytochrome P450 regulator Dap1 |
| *cbr1* | *SPCC970.03* | NADH-dependent reductase for Dph3, Cbr1 (predicted) |
| *ptl3* | *SPAC1A6.05c* | triacylglycerol lipase ptl3 |
|  | *SPBC215.11c* | aldo/keto reductase, unknown biological role |

<sup>a</sup> Descriptions were obtained from the *S. pombe* GeneDB (https://www.pombase.org).

### Ofd1-dependent regulation of SREBP may be conserved in other fungi

SREBP transcription factors are synthesized as inactive precursors that contain two predicted transmembrane segments [45]. The active SREBP N-terminal transcription factor domain (SREBP-N) is released by proteolytic cleavage and is regulated by Ofd1 in fission yeast [8]. Many fungal genomes contain putative SREBP or SREBP-N proteins and also code for Ofd1 homologs, but the role of these Ofd1 homologs in hypoxic adaptation is unknown [9]. Ofd1 negatively regulates cleaved Sre1N in fission yeast through direct binding to the bHLH domain of Sre1N [14]. To test whether Ofd1 may regulate SREBPs in other fungi, we performed yeast two-hybrid assays between fission yeast Ofd1 and the bHLH domains of 15 SREBPs or SREBP-N proteins from different fungi, including Sre1 and Sre2 from *S. pombe*. The majority of the tested bHLH domains (12/15) were expressed well as fusions to the Gal4 activation domain (Fig S7). Interestingly, fission yeast Ofd1 interacted strongly with 4 of the 12 bHLH domains from other fungal species: Hms1 and Tye7 of *S. cerevisae*, Tye7 of *C. albicans*, and *SRE1* (CNJ02310) in *C. neoformans* (Fig 4A, 4B). Hms1 and Tye7 are SREBP-N proteins that lack the transmembrane segments, suggesting that Ofd1 can regulate SREBP-N proteins in addition to proteolytic product of SREBP [9]. Each of these organisms contains an Ofd1 homolog with 35%-42% identity to *S. pombe* Ofd1 (Fig S8). Taken together, these results suggest that the regulation of SREBPs by Ofd1 may be conserved in other fungi.

**Fig 4.**
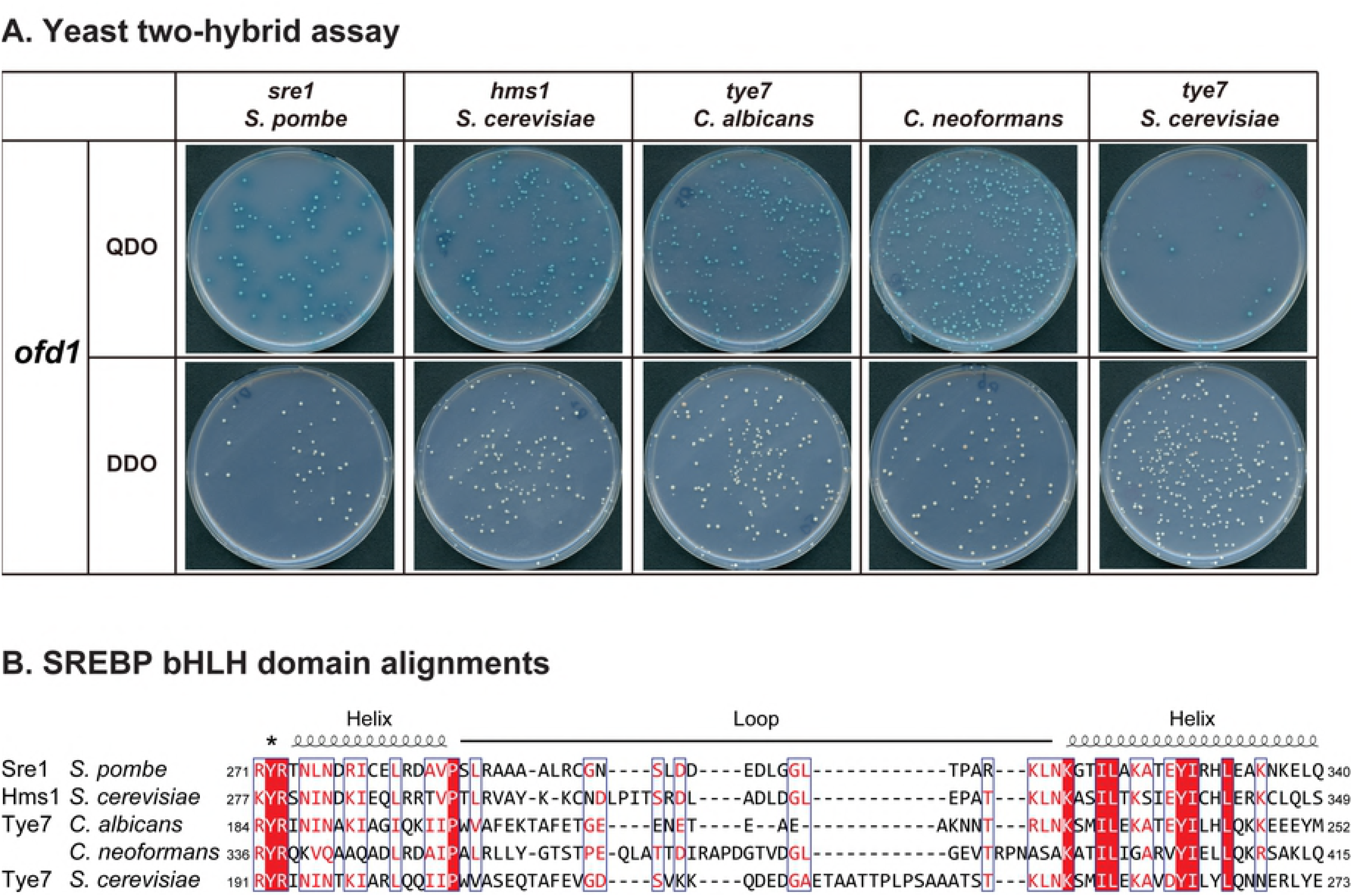
Regulation of SREBP transcription factor through Ofd1 may be conserved in other fungi. (A) Yeast two-hybrid interactions between basic helix-loop-helix (bHLH) regions of fungal SREBP or SREBP-N proteins and *S. pombe* Ofd1. bHLH regions of SREBPs were fused with GAL4 activation binding domain and Ofd1 was fused with GAL4 DNA binding domain. DDO: control plate, SD –Leu –Trp. QDO: selection plate, SD –Leu –Trp –Ade –His, supplemented with X-α-Gal. (B) Alignment of bHLH regions of SREBPs was performed by T-coffee. White character in red box: strict identity. Red character: similarity in one group. Blue frame: similarity across groups. ^∗^: tyrosine residue conserved in all SREBPs [34, 35].

## Discussion

In this study, we report three major findings: (1) novel binding partners and candidate prolyl hydroxylated substrates for the conserved 2-OG oxygenase Ofd1, (2) datasets of prolyl hydroxylated proteins in fission yeast eIF3m prep and the cellular proteome, and (3) that the principal impact of Ofd1 on the fission yeast proteome is through the regulation of SREBP, which may be conserved in other fungi.

We identified six novel Ofd1 binding partners by yeast two-hybrid screen (Fig 1) including two subunits of the eIF3 complex: Tif35 (eIF3g) and Sum1 (eIF3i). The other four binding partners are interesting proteins as well. Fin1 is a NIMA-related serine/threonine protein kinase, whose activity is regulated by Wee1 and Cdc25 and is required for G2/M transition [46, 47]. Isa2 is a mitochondrial iron-sulfur cluster assembly protein [48]. Pdb1 is the β subunit of pyruvate dehydrogenase E1[49]. Rps20 is a small ribosomal protein. Enrichment and targeted mass spectrometry studies are required to determine whether these binding partners are Ofd1 substrates.

In the mass spectrometry analysis of the eIF3m prep, the coverage of Tif35 and Sum1 was 63% and 58% respectively. However, we did not find Ofd1-dependent hydroxylations on these two proteins, suggesting that either the modified peptides are difficult to detect or that these proteins are Ofd1 binding partners but not enzymatic substrates. Indeed, Nro1 and Sre1 are Ofd1 binding partners, and no hydroxylated prolines have been found on these two proteins. Instead, we found 16 Ofd1 candidate substrates among eIF3m co-purifying proteins and provide strong evidence for Rpb10 as a substrate because the non-hydroxylated peptide accumulated in *ofd1Δ* cells (Fig 2, S1, Table 1). Rpb10 is an essential small subunit shared by RNA polymerases (RNAP) I, II and III, where it functions in the assembly of RNAP [50]. Rpb10 is composed of a N-terminal zinc bundle and a C-terminal tail, and Rpb10 directly interacts with Rpb2 and Rpb3 in RNAP II [51]. The hydroxylated proline in Rpb10, P64, lies toward the end of the C-terminal tail, and the proline residue is conserved in budding yeast and humans. Interestingly, Ofd1 prolyl hydroxylation of Rpb10 is quite different in two ways from that of Rps23. First, Rps23 is prolyl dihydroxylated, while Rpb10 is the first prolyl mono-hydroxylated substrate of Ofd1. Recently, we demonstrated that Nro1, a nuclear import adaptor, stabilizes the interaction between Ofd1 and its substrate Rps23, thus facilitating the dihydroxylation of Rps23 [17]. In *nro1Δ* cells, monohydroxylated Rps23 accumulated and dihydroxylated Rps23 decreased, suggesting stepwise hydroxylation reactions that depend on the strength of Ofd1-substrate interaction. Based on this, we predict that the interaction between Ofd1 and Rpb10 is weaker than that of Ofd1-Rps23-Nro1, which leads to the release of monohydroxylated Rpb10. Ofd1 dihydroxylates Rps23 P62 on the 3 and 4 positions, while the human homolog OGFOD1 is a prolyl 3-hydroxylase [18]. Structural studies of Ofd1 and its substrates Rpb10 and Rps23 will provide insight into the stereochemistry of these hydroxylation events. Second, Rps23 is completely hydroxylated, while there are measurable amounts of both non-hydroxylated and hydroxylated Rpb10 under normoxia. The existence of two populations of Rpb10 suggests that prolyl hydroxylation may play a regulatory role, making studies about the function of Rpb10 hydroxylation intriguing. Indeed, Rpb10 is not the only prolyl hydroxylated subunit in RNAP. Studies in human renal clear cell carcinoma cells suggested that the large subunit of RNAP II, Rpb1, is prolyl hydroxylated by PHD1 under low-grade oxidative-stress, which is necessary for the subsequent phosphorylation of Rpb1 and its non-degradative ubiquitination by VHL [52]. Whether Rpb10 hydroxylation is conserved in humans and whether there is a functional connection between the two prolyl hydroxylation events are interesting open questions.

Fission yeast Sre1 and its active form Sre1N regulate adaptation to low oxygen. Interaction between Ofd1 and Sre1N accelerates Sre1N degradation. Under hypoxia, this interaction is diminished and Sre1N accumulates, as Ofd1 is sequestered by binding to the Rps23-Nro1 complex [13, 14, 16, 17]. Although a hydroxylated proline on Sre1N has not been identified, this oxygen-dependent regulation of a transcription factor by a prolyl hydroxylase resembles the HIF pathway, where PHD prolyl hydroxylates HIFα subunit and thereby targets it for proteasomal degradation [2]. Our proteomic analysis of *WT* and *ofd1Δ* cells revealed that Ofd1 does not broadly regulate protein degradation. Instead, we observed an accumulation of proteins in *ofd1Δ* cells encoded by Sre1 target genes, indicating that the effect of Ofd1 on the proteome is through negative regulation of Sre1N (Fig 3, Table5).

To explore whether Ofd1 regulation of Sre1N exists in other fungal organisms, we tested whether this binding interaction is conserved. Yeast two-hybrid analysis between Ofd1 and the bHLH domains of fungal SREBP or SREBP-N proteins showed that four SREBP and SREBP-N proteins from *S. cerevisiae*, *C. albicans*, and *C. neoformans* interacted with Ofd1, suggesting a conserved regulation of SREBPs by Ofd1 in these species (Fig 4). We recently identified an Ofd1-binding motif in Sre1N aa 286-297, which is shared in Rps23 and Nro1 [17]. However, this motif is not apparent in the four interacting bHLH domains. Sequence alignments of Ofd1 homologs from these organisms showed a strong conservation of the catalytic N-terminal domain while the C-terminal domain (Fig S8) is less conserved. Given that the Ofd1 C-terminal domain binds Sre1N and mediates its degradation [13, 14], the C-terminal domains of the Ofd1 homologs may have co-evolved with the SREBP and SREBP-N proteins to preserve this regulatory protein-protein interaction.

Prolyl hydroxylation plays different roles in cells, ranging from a structural function in collagen biogenesis to regulation of the hypoxic response. In yeast, Rps23 is the only known prolyl hydroxylated protein, and hydroxylation functions to regulate Sre1N and promotes translational fidelity [17, 18]. In this study, we performed the first search for prolyl hydroxylation in fission yeast, and we discovered 12 additional prolyl hydroxylated proteins with complete mass spectrometry fragmentation around the hydroxylated proline. Even without an enrichment method, 4.2% of proteins were prolyl hydroxylated in our proteomic search, suggesting that prolyl hydroxylation is more prevalent than previously known. The development of a robust enrichment method is critical to fully reveal the prolyl hydroxylome.

The prolyl hydroxylation of Rps23 is conserved in many species and affects translation termination [17-20]. To date, three other ribosomal proteins have been found to be hydroxylated by 2-OG/Fe(II)-dependent oxygenases – arginine hydroxylation of *E. coli* Rpl16 by YcfD, histidine hydroxylation of human Rpl8 by NO66, and histidine hydroxylation of human Rpl27a by NO66 [53]. The functional role of these hydroxylations has yet to be described. Here, we found three additional ribosomal subunits with prolyl hydroxylations, Rps2, Rps9 and Rpl10a. Together, these findings suggest that the ribosome is a common target for protein hydroxylation. Further characterization of these modifications and their corresponding oxygenases will provide new insights into translation and cell proliferation.

## Data availability

The eIF3m prep SILAC mass spectrometry data and fission yeast whole proteome TMT mass spectrometry data have been deposited to the ProteomeXchange Consortium via the PRIDE partner repository with the dataset identifier PXD009372 and PXD009371, respectively [54].

## Acknowledgements

We thank the members of the Espenshade lab for their advice and acknowledge Bob O’Meally and Robert Cole from the JHMI Mass Spectrometry Core for analyzing the SILAC and TMT samples.

## Supporting information

**S1 Fig. Supplemental figures of Fig 2.** (A) Incorporation efficiency of heavy lysine in SILAC culture from Figure 2. The heavy lysate of *WT* cells was analyzed by mass spectrometry. Incorporation level was defined as the ratio of heavy lysine to total. Incorporation level was plotted using density plot in R. The median level of incorporation was 0.80 and this value was used for normalization. (B) Multiple sequence alignment of prospective Ofd1 binding sites in Pic1 and Fas1 was generated using T-coffee [34, 35]. (C) Comparison of prolyl hydroxylated and unmodified peptide from Rps23 and Rpb10 in eIF3m prep.

**S2 Fig. Ofd1 candidate substrates from eIF3m prep.** Spectra of Ofd1-dependent prolyl hydroxylated PSMs. Peptide sequences and fragment spectra are shown. PSMs with H/L > 3.5 were classified as Ofd1-dependent prolyl hydroxylations.

**S3 Fig. Ofd1-independent prolyl hydroxylations from eIF3m prep.** Spectra of Ofd1-independent prolyl-hydroxylated PSMs. Peptide sequences and fragment spectra are shown. PSMs with H/L ≤ 3.5 were classified as Ofd1-independent prolyl hydroxylations.

**S4 Fig. Analysis of the left-shifted population.** (A) Abundance of all peptides identified from eIF3m prep. Each PSM was plotted as a dot. The relative abundance of each peptide in *WT* and *ofd1*Δ cells was quantified by SILAC. Yellow: prolyl hydroxylated PSMs. Grey: all other PSMs. Distribution of H/L was plotted as histogram on the right. The small peak on the left corresponded to the left-shifted population. (B) Abundance of all proteins identified from eIF3m prep. Protein quantification was calculated by Proteome Discoverer 1.4, and was displayed as the average area of the three unique peptides with the largest peak area in heavy and light quantification channels. Each protein was plotted as a dot. Yellow: prolyl-hydroxylated PSMs. Grey: all other PSMs. (C) Protein half-life analysis. Protein half-life was obtained from published database [42], and was plotted as histogram using R. The distribution of half-life of all proteins identified in eIF3m prep (median = 12.9), left-shifted peptides (median = 13.0), prolyl hydroxylated peptides (median = 13.3, and left-shifted prolyl hydroxylated peptides (median = 13.4) were compared. Student t-tests showed no significant difference between the distributions.

**S5 Fig. Quantile normalization of TMT mass spectrometry data for analysis of WT and *ofd1Δ* proteomes.** The relative abundance of *WT5* and ofd1Δ2 was higher than the other replicates for unknown reasons, and therefore these two samples were excluded from quantile normalization and further analysis.

**S6 Fig. Ofd1-independent prolyl hydroxylations from *S. pombe* proteome.** Spectra of Ofd1-independent prolyl-hydroxylated PSMs. Peptide sequences and fragment spectra are shown.

**S7 Fig. Expression of fungal bHLH domains in Gal4-AD vector.** The bHLH domain of 15 SREBPs or SREBP-N proteins were cloned into Gal4 activation domain vector (pGADT7). HA tag was fused between Gal4 activation domain and the inserted bHLH domains and was used to detect the expression fusion proteins in yeast extracts by western blot. Asterisks denoted those fusion constructs tested in Figure 4. A non-specific band was used as loading control.

**S8 Fig. Sequence alignments of Ofd1 homologs.** Sequences from NCBI database were aligned using T-coffee [34, 35], and displayed with ESPript 3 [37]. The structure of Tpa1 from *S. cerevisiae* was used to predict the secondary structure indicated above the alignment. α-helices (α) and 3_10_-helices (η) are displayed as loops. β-strands are rendered as arrows, strict β-turns as TT letters and strict α-turns as TTT [36].

## References

1. Ploumakis A, Coleman ML. OH, the places you’ll Go! Hydroxylation, gene expression, and cancer. Mol Cell. 2015;58(5):729–41. doi: 10.1016/j.molcel.2015.05.026. PubMed PMID: 26046647.

2. Semenza GL. Regulation of cancer cell metabolism by hypoxia-inducible factor 1. Semin Cancer Biol. 2009;19(1):12–6. Epub 2008/12/31. doi: 10.1016/j.semcancer.2008.11.009. PubMed PMID: 19114105.

3. Ozer A, Bruick RK. Non-heme dioxygenases: cellular sensors and regulators jelly rolled into one? Nat Chem Biol. 2007;3(3):144–53. doi: 10.1038/nchembio863. PubMed PMID: 17301803.

4. Kaelin WG, Jr., Ratcliffe PJ. Oxygen sensing by metazoans: the central role of the HIF hydroxylase pathway. Mol Cell. 2008;30(4):393–402. Epub 2008/05/24. doi: 10.1016/j.molcel.2008.04.009. PubMed PMID: 18498744.

5. Zurlo G, Guo J, Takada M, Wei W, Zhang Q. New insights into protein hydroxylation and its important role in human diseases. Biochim Biophys Acta. 2016;1866(2):208–20. Epub 2016/10/25. doi: 10.1016/j.bbcan.2016.09.004. PubMed PMID: 27663420; PubMed Central PMCID: PMCPMC5138100.

6. Semenza GL. Targeting HIF-1 for cancer therapy. Nat Rev Cancer. 2003;3(10):721–32. Epub 2003/09/18. doi: 10.1038/nrc1187. PubMed PMID: 13130303.

7. Semenza GL. Hypoxia-inducible factors in physiology and medicine. Cell. 2012;148(3):399–408. doi: 10.1016/j.cell.2012.01.021. PubMed PMID: WOS:000300225000010.

8. Hughes AL, Todd BL, Espenshade PJ. SREBP pathway responds to sterols and functions as an oxygen sensor in fission yeast. Cell. 2005;120(6):831–42. doi: 10.1016/j.cell.2005.01.012. PubMed PMID: 15797383.

9. Bien CM, Espenshade PJ. Sterol regulatory element binding proteins in fungi: hypoxic transcription factors linked to pathogenesis. Eukaryot Cell. 2010;9(3):352–9. doi: 10.1128/EC.00358-09. PubMed PMID: 20118213; PubMed Central PMCID: PMCPMC2837984.

10. Chang YC, Bien CM, Lee H, Espenshade PJ, Kwon-Chung KJ. Sre1p, a regulator of oxygen sensing and sterol homeostasis, is required for virulence in *Cryptococcus neoformans*. Molecular Microbiology. 2007;64(3):614–29. doi: 10.1111/j.1365-2958.2007.05676.x. PubMed PMID: WOS:000245991000005.

11. Willger SD, Puttikamonkul S, Kim KH, Burritt JB, Grahl N, Metzler LJ, et al. A sterol-regulatory element binding protein is required for cell polarity, hypoxia adaptation, azole drug resistance, and virulence in *Aspergillus fumigatus*. Plos Pathog. 2008;4(11). doi: 10.1371/journal.ppat.1000200. PubMed PMID: WOS:000261481200005.

12. Maguire SL, Wang C, Holland LM, Brunel F, Neuvéglise C, Nicaud JM, et al. Zinc finger transcription factors displaced SREBP proteins as the major Sterol regulators during *Saccharomycotina* evolution. PLoS Genet. 2014;10(1):e1004076. doi: 10.1371/journal.pgen.1004076. PubMed PMID: 24453983; PubMed Central PMCID: PMCPMC3894159.

13. Hughes BT, Espenshade PJ. Oxygen-regulated degradation of fission yeast SREBP by Ofd1, a prolyl hydroxylase family member. EMBO J. 2008;27(10):1491–501. doi: 10.1038/emboj.2008.83. PubMed PMID: 18418381; PubMed Central PMCID: PMCPMC2396400.

14. Lee CY, Yeh TL, Hughes BT, Espenshade PJ. Regulation of the Sre1 hypoxic transcription factor by oxygen-dependent control of DNA binding. Mol Cell. 2011;44(2):225–34. doi: 10.1016/j.molcel.2011.08.031. PubMed PMID: 22017871; PubMed Central PMCID: PMCPMC3208185.

15. Porter JR, Lee CY, Espenshade PJ, Iglesias PA. Regulation of SREBP during hypoxia requires Ofd1-mediated control of both DNA binding and degradation. Mol Biol Cell. 2012;23(18):3764–74. doi: 10.1091/mbc.E12-06-0451. PubMed PMID: 22833559; PubMed Central PMCID: PMCPMC3442422.

16. Lee CY, Stewart EV, Hughes BT, Espenshade PJ. Oxygen-dependent binding of Nro1 to the prolyl hydroxylase Ofd1 regulates SREBP degradation in yeast. EMBO J. 2009;28(2):135–43. doi: 10.1038/emboj.2008.271. PubMed PMID: 19158663; PubMed Central PMCID: PMCPMC2634736.

17. Clasen SJ, Shao W, Gu H, Espenshade PJ. Prolyl dihydroxylation of unassembled uS12/Rps23 regulates fungal hypoxic adaptation. Elife. 2017;6. Epub 2017/10/31. doi: 10.7554/eLife.28563. PubMed PMID: 29083304; PubMed Central PMCID: PMCPMC5690285.

18. Loenarz C, Sekirnik R, Thalhammer A, Ge W, Spivakovsky E, Mackeen MM, et al. Hydroxylation of the eukaryotic ribosomal decoding center affects translational accuracy. Proc Natl Acad Sci U S A. 2014;111(11):4019–24. doi: 10.1073/pnas.1311750111. PubMed PMID: 24550462; PubMed Central PMCID: PMCPMC3964080.

19. Singleton RS, Liu-Yi P, Formenti F, Ge W, Sekirnik R, Fischer R, et al. OGFOD1 catalyzes prolyl hydroxylation of RPS23 and is involved in translation control and stress granule formation. Proc Natl Acad Sci U S A. 2014;111(11):4031–6. doi: 10.1073/pnas.1314482111. PubMed PMID: 24550447; PubMed Central PMCID: PMCPMC3964040.

20. Katz MJ, Acevedo JM, Loenarz C, Galagovsky D, Liu-Yi P, Pérez-Pepe M, et al. Sudestada1, a Drosophila ribosomal prolyl-hydroxylase required for mRNA translation, cell homeostasis, and organ growth. Proc Natl Acad Sci U S A. 2014;111(11):4025–30. doi: 10.1073/pnas.1314485111. PubMed PMID: 24550463; PubMed Central PMCID: PMCPMC3964085.

21. Hutton JJ, Jr., Kaplan A, Udenfriend S. Conversion of the amino acid sequence gly-pro-pro in protein to gly-pro-hyp by collagen proline hydroxylase. Arch Biochem Biophys. 1967;121(2):384–91. Epub 1967/08/01. PubMed PMID: 6057106.

22. Kivirikko KI, Prockop DJ. Enzymatic hydroxylation of proline and lysine in protocollagen. Proc Natl Acad Sci U S A. 1967;57(3):782–9. Epub 1967/03/01. PubMed PMID: 16591531; PubMed Central PMCID: PMCPMC335576.

23. Wong BW, Kuchnio A, Bruning U, Carmeliet P. Emerging novel functions of the oxygen-sensing prolyl hydroxylase domain enzymes. Trends Biochem Sci. 2013;38(1):3–11. doi: 10.1016/j.tibs.2012.10.004. PubMed PMID: 23200187.

24. Xie L, Xiao K, Whalen EJ, Forrester MT, Freeman RS, Fong G, et al. Oxygen-regulated beta(2)-adrenergic receptor hydroxylation by EGLN3 and ubiquitylation by pVHL. Sci Signal. 2009;2(78):ra33. Epub 2009/07/09. doi: 10.1126/scisignal.2000444. PubMed PMID: 19584355; PubMed Central PMCID: PMCPMC2788937.

25. Heir P, Srikumar T, Bikopoulos G, Bunda S, Poon BP, Lee JE, et al. Oxygen-dependent regulation of erythropoietin receptor turnover and signaling. J Biol Chem. 2016;291(14):7357–72. Epub 2016/02/06. doi: 10.1074/jbc.M115.694562. PubMed PMID: 26846855; PubMed Central PMCID: PMCPMC4817168.

26. Luo WB, Hu HX, Chang R, Zhong J, Knabel M, O’Meally R, et al. Pyruvate kinase M2 is a PHD3-stimulated coactivator for hypoxia-inducible factor 1. Cell. 2011;145(5):732–44. doi: 10.1016/j.cell.20n.03.054. PubMed PMID: WOS:000291018600013.

27. Koditz J, Nesper J, Wottawa M, Stiehl DP, Camenisch G, Franke C, et al. Oxygen-dependent ATF-4 stability is mediated by the PHD3 oxygen sensor. Blood. 2007;110(10):3610–7. doi: DOI 10.1182/blood-2007-06-094441. PubMed PMID: WOS:000250946300026.

28. Bahler J, Wu JQ, Longtine MS, Shah NG, McKenzie A, 3rd, Steever AB, et al. Heterologous modules for efficient and versatile PCR-based gene targeting in *Schizosaccharomyces pombe*. Yeast. 1998;14(10):943–51. Epub 1998/08/26. doi: 10.1002/(SICI)1097-0061(199807)14:10<943::AID-YEA292>3.0.CO;2-Y. PubMed PMID: 9717240.

29. Fröhlich F, Christiano R, Walther TC. Native SILAC: metabolic labeling of proteins in prototroph microorganisms based on lysine synthesis regulation. Mol Cell Proteomics. 2013;12(7):1995–2005. doi: 10.1074/mcp.M112.025742. PubMed PMID: 23592334; PubMed Central PMCID: PMCPMC3708181.

30. James P, Halladay J, Craig EA. Genomic libraries and a host strain designed for highly efficient two-hybrid selection in yeast. Genetics. 1996;144(4):1425–36. Epub 1996/12/01. PubMed PMID: 8978031; PubMed Central PMCID: PMCPMC1207695.

31. Hwang J, Ribbens D, Raychaudhuri S, Cairns L, Gu H, Frost A, et al. A Golgi rhomboid protease Rbd2 recruits Cdc48 to cleave yeast SREBP. EMBO J. 2016;35(21):2332–49. doi: 10.15252/embj.201693923. PubMed PMID: 27655872; PubMed Central PMCID: PMCPMC5090219.

32. Bolstad B. preprocessCore: A collection of pre-processing functions. R package version 1.40.0. https://github.com/bmbolstad/preprocessCore. 2017.

33. Stewart EV, Nwosu CC, Tong Z, Roguev A, Cummins TD, Kim DU, et al. Yeast SREBP cleavage activation requires the Golgi Dsc E3 ligase complex. Mol Cell. 2011;42(2):160–71. doi: 10.1016/j.molcel.2011.02.035. PubMed PMID: 21504829; PubMed Central PMCID: PMCPMC3083633.

34. Notredame C, Higgins DG, Heringa J. T-Coffee: A novel method for fast and accurate multiple sequence alignment. J Mol Biol. 2000;302(1):205–17. doi: 10.1006/jmbi.2000.4042. PubMed PMID: 10964570.

35. Di Tommaso P, Moretti S, Xenarios I, Orobitg M, Montanyola A, Chang JM, et al. T-Coffee: a web server for the multiple sequence alignment of protein and RNA sequences using structural information and homology extension. Nucleic Acids Res. 2011;39(Web Server issue):W13–7. doi: 10.1093/nar/gkr245. PubMed PMID: 21558174; PubMed Central PMCID: PMCPMC3125728.

36. Kim HS, Kim HL, Kim KH, Kim DJ, Lee SJ, Yoon JY, et al. Crystal structure of Tpa1 from *Saccharomyces cerevisiae*, a component of the messenger ribonucleoprotein complex. Nucleic Acids Research. 2010;38(6):2099–110. doi: 10.1093/nar/gkp1151. PubMed PMID: WOS:000276304600038.

37. Robert X, Gouet P. Deciphering key features in protein structures with the new ENDscript server. Nucleic Acids Res. 2014;42(Web Server issue):W320–4. doi: 10.1093/nar/gku316. PubMed PMID: 24753421; PubMed Central PMCID: PMCPMC4086106.

38. Zhou C, Arslan F, Wee S, Krishnan S, Ivanov AR, Oliva A, et al. PCI proteins eIF3e and eIF3m define distinct translation initiation factor 3 complexes. BMC Biol. 2005;3:14. doi: 10.1186/1741-7007-3-14. PubMed PMID: 15904532; PubMed Central PMCID: PMCPMC1173091.

39. Sha Z, Brill LM, Cabrera R, Kleifeld O, Scheliga JS, Glickman MH, et al. The eIF3 interactome reveals the translasome, a supercomplex linking protein synthesis and degradation machineries. Mol Cell. 2009;36(1):141–52. Epub 2009/10/13. doi: 10.1016/j.molcel.2009.09.026. PubMed PMID: 19818717; PubMed Central PMCID: PMCPMC2789680.

40. Hughes AL, Powell DW, Bard M, Eckstein J, Barbuch R, Link AJ, et al. Dap1/PGRMC1 binds and regulates cytochrome P450 enzymes. Cell Metab. 2007;5(2):143–9. Epub 2007/02/06. doi: 10.1016/j.cmet.2006.12.009. PubMed PMID: 17276356.

41. Rigaut G, Shevchenko A, Rutz B, Wilm M, Mann M, Seraphin B. A generic protein purification method for protein complex characterization and proteome exploration. Nat Biotechnol. 1999;17(10):1030–2. Epub 1999/10/03. doi: 10.1038/13732. PubMed PMID: 10504710.

42. Christiano R, Nagaraj N, Frohlich F, Walther TC. Global proteome turnover analyses of the Yeasts *S. cerevisiae* and *S. pombe*. Cell Rep. 2014;9(5):1959–65. Epub 2014/12/04. doi: 10.1016/j.celrep.2014.10.065. PubMed PMID: 25466257; PubMed Central PMCID: PMCPMC4526151.

43. Wilhelm BT, Marguerat S, Watt S, Schubert F, Wood V, Goodhead I, et al. Dynamic repertoire of a eukaryotic transcriptome surveyed at single-nucleotide resolution. Nature. 2008;453(7199):1239–43. doi: 10.1038/nature07002. PubMed PMID: 18488015.

44. Todd BL, Stewart EV, Burg JS, Hughes AL, Espenshade PJ. Sterol regulatory element binding protein is a principal regulator of anaerobic gene expression in fission yeast. Mol Cell Biol. 2006;26(7):2817–31. doi: 10.1128/MCB.26.7.2817-2831.2006. PubMed PMID: 16537923; PubMed Central PMCID: PMCPMC1430309.

45. Espenshade PJ, Hughes AL. Regulation of sterol synthesis in eukaryotes. Annu Rev Genet. 2007;41:401–27. Epub 2007/08/02. doi: 10.1146/annurev.genet.41.110306.130315. PubMed PMID: 17666007.

46. Grallert A, Connolly Y, Smith DL, Simanis V, Hagan IM. The *S. pombe* cytokinesis NDR kinase Sid2 activates Fin1 NIMA kinase to control mitotic commitment through Pom1/Wee1. Nat Cell Biol. 2012;14(7):738–45. Epub 2012/06/12. doi: 10.1038/ncb2514. PubMed PMID: 22684255; PubMed Central PMCID: PMCPMC4284365.

47. Krien MJ, Bugg SJ, Palatsides M, Asouline G, Morimyo M, O’Connell MJ. A NIMA homologue promotes chromatin condensation in fission yeast. J Cell Sci. 1998;111 (Pt 7):967–76. Epub 1998/05/20. PubMed PMID: 9490640.

48. Kim KD, Chung WH, Kim HJ, Lee KC, Roe JH. Monothiol glutaredoxin Grx5 interacts with FeS scaffold proteins Isa1 and Isa2 and supports Fe-S assembly and DNA integrity in mitochondria of fission yeast. Biochem Biophys Res Commun. 2010;392(3):467–72. Epub 2010/01/21. doi: 10.1016/j.bbrc.2010.01.051. PubMed PMID: 20085751.

49. Cavan G, MacDonald D. Cloning and sequence of a gene encoding the pyruvate dehydrogenase E1 beta subunit of *Schizosaccharomyces pombe*. Gene. 1995;152(1):117–20. Epub 1995/01/11. PubMed PMID: 7828917.

50. Woychik NA, Young RA. RNA polymerase-II subunit Rpb10 is essential for yeast cell viability. Journal of Biological Chemistry. 1990;265(29):17816–9. PubMed PMID: WOS:A1990EC76600070.

51. Cramer P, Bushnell DA, Kornberg RD. Structural basis of transcription: RNA polymerase II at 2.8 angstrom ngstrom resolution. Science. 2001;292(5523):1863–76. doi: DOI 10.1126/science.1059493. PubMed PMID: WOS:000169200700039.

52. Mikhaylova O, Ignacak ML, Barankiewicz TJ, Harbaugh SV, Yi Y, Maxwell PH, et al. The von Hippel-Lindau tumor suppressor protein and Egl-9-Type proline hydroxylases regulate the large subunit of RNA polymerase II in response to oxidative stress. Mol Cell Biol. 2008;28(8):2701–17. Epub 2008/02/21. doi: 10.1128/MCB.01231-07. PubMed PMID: 18285459; PubMed Central PMCID: PMCPMC2293119.

53. Ge W, Wolf A, Feng T, Ho CH, Sekirnik R, Zayer A, et al. Oxygenase-catalyzed ribosome hydroxylation occurs in prokaryotes and humans. Nat Chem Biol. 2012;8(12):960–2. doi: 10.1038/nchembio.1093. PubMed PMID: 23103944; PubMed Central PMCID: PMCPMC4972389.

54. Vizcaino JA, Csordas A, Del-Toro N, Dianes JA, Griss J, Lavidas I, et al. 2016 update of the PRIDE database and its related tools. Nucleic Acids Res. 2016;44(22):11033. Epub 2016/09/30. doi: 10.1093/nar/gkw880. PubMed PMID: 27683222; PubMed Central PMCID: PMCPMC5159556.

